# Developmental pathways underlying sexual differentiation in a U/V sex chromosome system

**DOI:** 10.1101/2024.02.09.579736

**Authors:** Daniel Liesner, Guillaume Cossard, Min Zheng, Olivier Godfroy, Josué Barrera-Redondo, Fabian B. Haas, Susana M. Coelho

## Abstract

In many multicellular organisms, sexual development is not determined by XX/XY or ZW/ZZ systems but by U/V sex chromosomes. In U/V systems, sex determination occurs in the haploid phase, with U chromosomes in females and V chromosomes in males. Here, we explore several male, female and partially sex-reversed male lines of giant kelp to decipher how U/V sex chromosomes and autosomes initiate male versus female development. We identify a key set of genes on the sex chromosomes involved in triggering sexual development, and characterise autosomal effector genes underlying sexual differentiation. We show that male, but not female, development involves large-scale transcriptome reorganisation with pervasive enrichment in regulatory genes, faster evolutionary rates, and high species specificity of male-biased genes. Our observations imply that a female-like phenotype is the “ground state”, which is complemented by the presence of a U-chromosome, but overridden by a dominant male developmental program in the presence of a V-chromosome.

## Introduction

In many eukaryotes, the establishment of the male versus the female developmental programs depends on a sex determining region (SDR) located on sex chromosomes^1,2^. Gene(s) carried in this region trigger the differential expression of a cascade of autosomal effector genetic pathways, eventually leading to sex-biased gene expression^3,4^. Sex determination and sexual differentiation have been extensively studied in animals and plants, often by genomic and transcriptomic comparison between sexes (see, *e.g.*, reviews by Charlesworth^5^; Feng et al.^6^) but also using sex-reversed mutants^7–10^. Beyond the causal mutation which often highlights key sex determining genes, the restructuring of autosomal gene expression in these mutants is expected to shed light on the core molecular pathways involved in sexual differentiation^11,12^. However, in groups of multicellular eukaryotes besides animals and some plants, knowledge on the key genes involved in sex phenotype differentiation remains elusive^13,14^.

Brown algae are one of few eukaryotic lineages to have independently evolved complex multicellularity^15^. Most brown algae are dioicous, *i.e.*, male and female sexes are determined in the haploid (gametophyte) stage^16^ via a U/V sexual system, in which females possess a U chromosome and males a V chromosome^17^. Sexual dimorphism of gametophytes is driven by sex-biased expression of autosomal genes^18^ operating downstream of master regulator(s) located on the sex-determining regions (SDR) of the U and V sex chromosomes^17,19^. The brown algal master male-determining gene (male inducer, *MIN*)^20^ located on the V-SDR has recently been identified, and encodes a HMG-box transcription factor^17,19–23^.

The filamentous gametophytes of the brown algal order Laminariales (*i.e.*, kelps) present a relatively simple sexual dimorphism^24^. Female gametophyte cells are larger, contain larger chloroplasts^24^, and upon gametogenesis develop into enlarged oogonia (**Figure 1A**). One immotile egg cell of 20–45 µm diameter^19,25^ is produced and remains attached to the oogonium^26^. Male gametophytes, in contrast, consist of less pigmented cells with roughly half the diameter of female cells^24^. Antheridia develop by mitosis on somatic cells and produce motile sperm of 5–9 µm length^19,27^. Sperm release and attraction is facilitated by the pheromone lamoxirene which is produced exclusively by the eggs^28^. Syngamy and embryo development lead to the complex multicellular diploid sporophyte generation, which may grow to tens of meters in length (**Figure 1A**). In mature sporophytes, meiosporangia are arranged in distinct sori, which in some species develop on specialised sporophylls, and release haploid meiospores, which settle and germinate into initial gametophyte cells (**Figure 1A**). Unfertilized eggs, but not sperm, may also develop parthenogenetically into partheno-sporophytes which are often deformed and not viable^29,30^.

**Figure 1.**
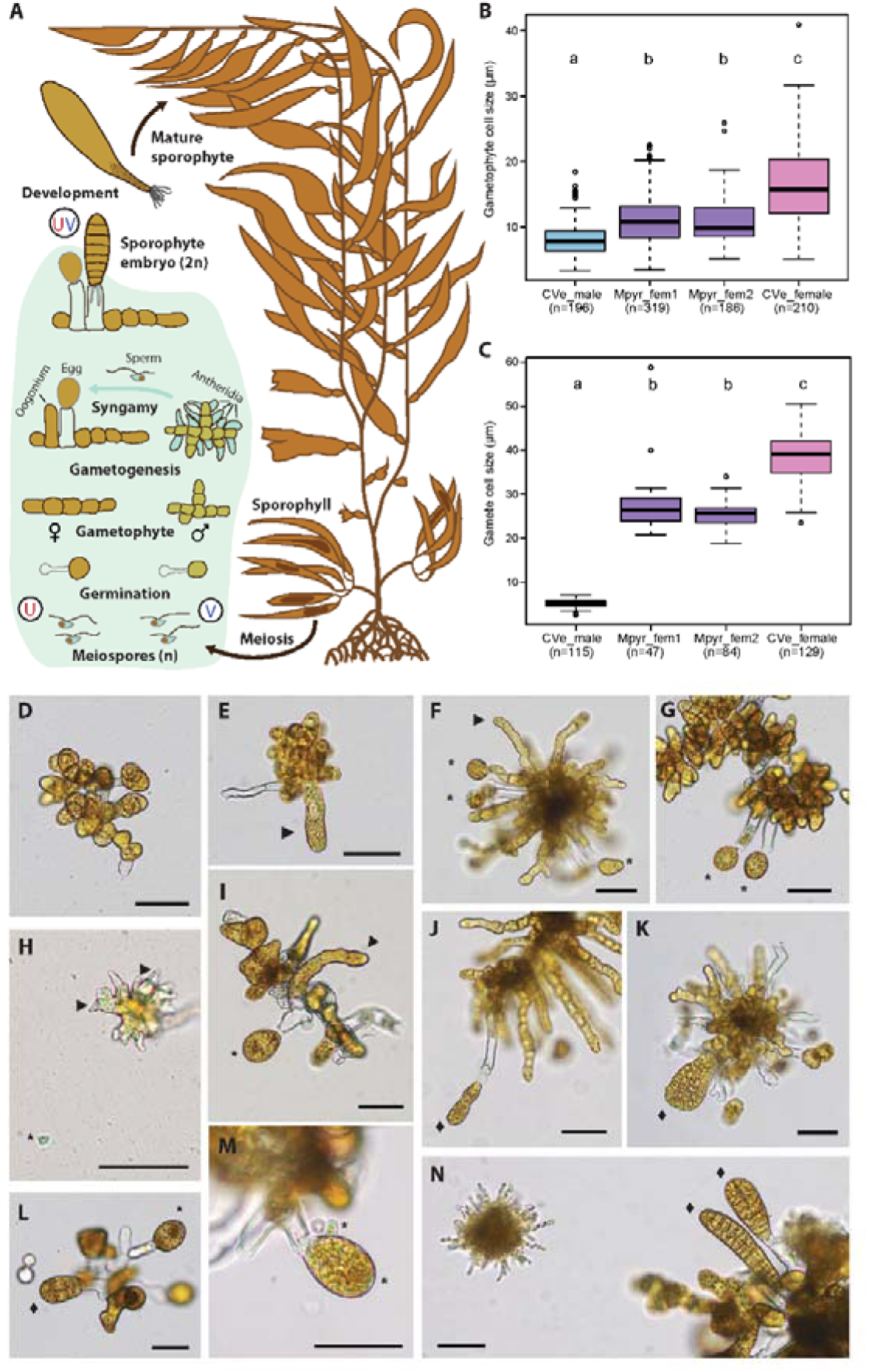
Morphology and observations of *Macrocystis pyrifera* WT male (CVe_male) and female (CVe_female) gametophytes, and the feminised male gametophytes Mpyr_fem1 and Mpyr_fem2. (A) Schematic life cycle of *M. pyrifera*, with microscopic, haploid stages highlighted in light green. Mature sporophytes (2n) release meiospores (n) from sori located on sporophylls. Meiospores settle and germinate into distinct, microscopic female and male gametophytes, which carry the U and V chromosome, respectively. Following gametogenesis, females release eggs from oogonia, which release the pheromone lamoxirene. This triggers the release and attraction of spermatozoids from male antheridia. Syngamy initiates the next sporophyte generation. Partheno-sporophytes may develop from unfertilised eggs (omitted for clarity). (B) Gametophyte cell size and (C) gamete cell size of WT male and female gametophytes and the feminised males. Letters above plot indicate significant differences between strains (Wilcoxon test, *p*<0.0001). (D) Vegetative Mpyr_fem2 gametophyte. (E) Mpyr_fem1 gametophyte with gametangium (arrow). (F) Mpyr_fem2 with gametangium (arrow) and released gametes (asterisks). (G) Mpyr_fem1 gametophyte with released gametes (asterisks). (H) Fertile WT male gametophyte with antheridia (arrows) and spermatozoid (asterisk). (I) Fertile WT female with oogonium (arrow) and released eggs (asterisk). (J) Mpyr_fem1 gametophyte with partheno-sporophyte (diamond). (K) Mpyr_fem2 gametophyte with partheno-sporophyte (diamond). (L) WT female gametophyte with partheno-sporophyte (diamond). (M) Fusion of male and female gametes (asterisks). (N) Sporophytes obtained from syngamy (diamonds) in co-cultivation of male and female gametophytes. All scale bars=50 µm.

Recently, a genetically male individual with a partially sex-reversed phenotype has been described in the giant kelp *Macrocystis pyrifera* ^19^. Males of the variant line Mpyr_13-4 exhibit an increased cell size intermediate to female and male wild type (WT) lines, and produce similarly enlarged, immotile gametes which are capable of parthenogenesis, therefore resembling eggs. However, these gametes do not produce the sperm-attracting pheromone lamoxirene^19^ and are therefore sterile. It was concluded that these individuals are only partially sex reversed, morphologically ‘feminised’. Accordingly, their transcriptome showed both an increased expression of female-biased genes (transcriptomic feminisation) and decreased expression of male-biased genes (transcriptomic de-masculinisation). Interestingly, two male SDR genes were significantly down-regulated in this line compared to the WT male, including the master male-determining gene *MIN*^19,20^. However, the sterility of the line precluded a detailed genetic analysis of the causative and effector genes underlying this phenotype.

Here, we exploit two additional, independently obtained giant kelp lines that are genetically male but present feminised developmental patterns, to examine the core molecular processes underlying sexual dimorphism in a haploid U/V sexual system. We estimate the extent of conservation of the molecular events underlying the male versus female developmental program, to disentangle the role of sex-linked genes and autosomal gene expression in the initiation of sex-specific development, and examine the conservation of the gene regulatory networks that underlie sexual dimorphism in these organisms. We show that feminisation is consistently associated with effector genes related to central metabolic pathways, as well as a subset of genes located on the sex chromosomes. Finally, our results suggest that male development involves large-scale restructuring of gene expression patterns and imply that a female-like phenotype is the “ground state” of the giant kelp morphology, which is complemented by the presence of a U-chromosome, but overridden by a dominant male developmental program in the presence of a V-chromosome.

## Results and discussion

### Identification of genetically male lines showing feminised developmental patterns

We had previously reported the identification of the genetically male giant kelp line Mpyr_13-4, which originated from a male gametophyte collected in Curahue (Cur), Chile, following treatment with colchicine^19^ (**Figure S1A**). Mpyr_13-4 presented a range of female features, but was sterile. We reasoned that comparing independently obtained male lines with a similarly feminised phenotype should allow to study the extent of conservation of the pathways involved in sexual differentiation and to pinpoint crucial genes required for male and female developmental functions. We identified two additional genetically male lines exhibiting a feminised phenotype, Mpyr_fem1 and Mpyr_fem2, originating from another population (Curaco de Vélez, Chile; CVe) that had been subjected to a colchicine treatment independently (**Figure 1**; **Figure S1**)**B**. PCR markers confirmed that these lines were *bona fide* genetically male (*i.e.*, had only the V and not the U-sex chromosome; **Figure S1C**). Flow cytometry analysis of DNA content provided no evidence for genome doubling compared to WT male samples (**Figure S1D**). Lack of aneuploidy was confirmed by Illumina sequencing of genomic DNA libraries. We mapped the reads to chromosome-level scaffolds of *M. pyrifera*^31^ and compared normalised sequence coverage per scaffold between wild type males and feminised variants (see methods). No scaffolds were identified as potentially duplicated or deleted using defined cut-offs and manual curation (**Figure S1E,F**). Therefore, despite the common use of colchicine as an inducer of polyploidy, none of the three variant lines showed signs of chromosome doubling (see also Müller et al.^19^). We conclude that Mpyr_fem1 and Mpyr_fem2 represent two independently feminised lines, in addition to Mpyr_13-4^19^.

A detailed morphometric analysis of these two variant lines showed that gametophyte cells of both variants were of intermediate size compared to the WT male and female lines (**Figure 1B**). Cell size differed significantly between lines, with variant cells being significantly smaller than WT female cells and significantly larger than WT male cells (**Figure 1B**; Kruskal-Wallis Χ²=283.64, df=3, *p<*0.0001).

During induction of gametogenesis (see methods), gametophyte cells of both variant lines developed into elongated gametangia, which resembled female oogonia (Figure 1D-F). From these, large, immotile gametes were released (Figure 1F-G). Gamete size differed significantly between lines (Figure 1C; Kruskal-Wallis Χ²=319.51, df=3, *p <*0.0001), with variant gametes (Figure 1F-G) being significantly larger than WT male gametes (Figure 1H), but significantly smaller than WT female gametes (Figure 1I). Note that gamete sizes did not differ significantly in a direct comparison between all three variants (Wilcoxon test; *p*_Mpyr_fem1-Mpyr_fem2=_0.084; *p*_Mpyr_fem1-Mpyr_13-4=_0.084; *p*_Mpyr_fem2-Mpyr_13-4=_0.144).

In *M. pyrifera*, only female gametes may undergo parthenogenesis if they are not fertilised by male gametes (Figure 1L), leading to development of partheno-sporophytes that morphologically resemble diploid sporophytes^29,32^. We observed development of partheno-sporophytes from unfertilised gametes in both genetically male variants (Figure 1J-K). We next tested whether the egg-like gametes were capable of attracting male gametes. When we brought fertile male gametophytes in contact with a WT female strain, sperm was immediately released, attraction and gamete fusion occurred (Figure 1M), and sporophytes developed within few days (Figure 1N). In stark contrast, Mpyr_fem1 and Mpyr_fem2 did not trigger sperm release and no gamete fusion occurred. These observations suggest that, similarly to the variant Mpyr_13-4 described previously^19^, Mpyr_fem1 and Mpyr_fem2 are incapable of attracting and fusing with males, suggesting the egg-like gametes do not produce the pheromone lamoxirene.

Together, these results suggest that although the two variant lines share morphological features with females (*e.g.*, larger cell dimensions, gamete morphology, parthenogenetic capacity), they are only partially sex-reversed, *i.e.*, they are not fully functional females.

### Transcriptomic patterns associated with sexual differentiation reveal high turnover of male-biased genes

In order to investigate the molecular pathways associated with the morphological feminisation of the variant lines, we used an RNA-seq approach using triplicate samples from the variants Mpyr_fem1 and Mpyr_fem2 in comparison to the WT male from which they were derived and to a WT female from the same population (CVe; **Table S1**; **Figure S**).**1**Our analysis also included the similarly feminised line Mpyr_13-4, which was independently obtained from a male of a different natural population (Cur)^19^, using publicly available datasets (**Table S1**; **Figure S1**).

Hierarchical clustering based on z-scores calculated from normalised expression values (transcripts per million, TPM) for all expressed genes (TPM > 5^th^ percentile; 18432 for CVe, 19301 for Cur) indicated similarity among replicates of the same strain (**Figure S2A,B)**, which was confirmed with principal component analyses (**Figure S2C,D**). Global gene expression patterns of all feminised variants Mpyr_fem1, Mpyr_fem2 and Mpyr_13-4 clustered intermediately to the respective WT males and females (**Figure S2A-D**). While the distinct cluster formed by Mpyr_fem1 and Mpyr_fem2 shared a node with the WT male CVe30m from which they arose, the feminised variant Mpyr_13-4^19^ clustered more closely with the WT female Cur4f than with the WT male Cur6m (**Figure S2A,B**). This result suggests that the global transcriptome of Mpyr_13-4 is slightly closer to that of a WT female, although the phenotypic changes are similar across all variants.

Male- and female-specific phenotypes are commonly driven by sex-biased expression of autosomal genes^18^. In order to further investigate the morphological feminisation of the variants, we focused on genes that were differentially expressed between males and females (*i.e.*, sex-biased genes). To account for batch effects between experiments (this study, Müller et al.^19^), we treated each population separately by extracting separate sets of differentially expressed genes (DEG) between WT males and WT females within each the CVe and Cur population using DESeq2^33^ (log_2_ fold-change, log_2_FC>1, padj<0.001; see methods).

For the CVe population, we found 1375 male-biased genes (MBG) and 1631 female-biased genes (FBG; **Table S2**). For the Cur population we found 2287 MBG and 2387 FBG (**Table S2**). 4926 genes were considered unbiased (log_2_FC<1, padj<0.001 in at least one population and unbiased or neither sex-biased nor unbiased in the other). The two populations shared 682 FBG and 489 MBG (**Figure S2E**) so only 24.1% of all FBG and 16.2% of all MBG were consistently sex biased in the same sex in both populations. Genetic heterogeneity among isolated gametophyte individuals^32^ may strongly influence sex-biased gene expression between these populations in our experiment, highlighting that potentially only a relatively small set of genes is driving sexual differentiation. Interestingly, we noticed that the fraction of shared FBG was significantly higher than that of shared MBG (test of equal proportions; Χ²=27.91, *p*=1.27*10^−7^), *i.e.*, MBG appear to show a higher variability between the two populations compared to FBG. We therefore investigated the turn-over of sex bias across larger phylogenetic distances (Figure 2A-B), using published datasets from four other brown algal species with separate sexes^34^. We found 1126 single-copy orthogroups which were expressed across all species, with 449 female-biased and 117 male-biased genes in at least one species (**Table S3**). Among these, we detected that subsets of female-biased orthologs of (at least one population of) *M. pyrifera* were also expressed as female-biased in all other species (Figure 2A). Female-biased orthologs were significantly enriched (fold-enrichment>1, *p*<0.001) among the shared orthologs of *M. pyrifera* with two other oogamous species, *Saccorhiza polyschides* and *Desmarestia herbacea* (multi-set intersection analysis^35^, Figure 2A). In stark contrast, no orthologs of MBG of *M. pyrifera* were male-biased in any of the four other species (Figure 2B). This pattern persists in *M. pyrifera* despite the low overlap of sex-biased genes between populations.

**Figure 2.**
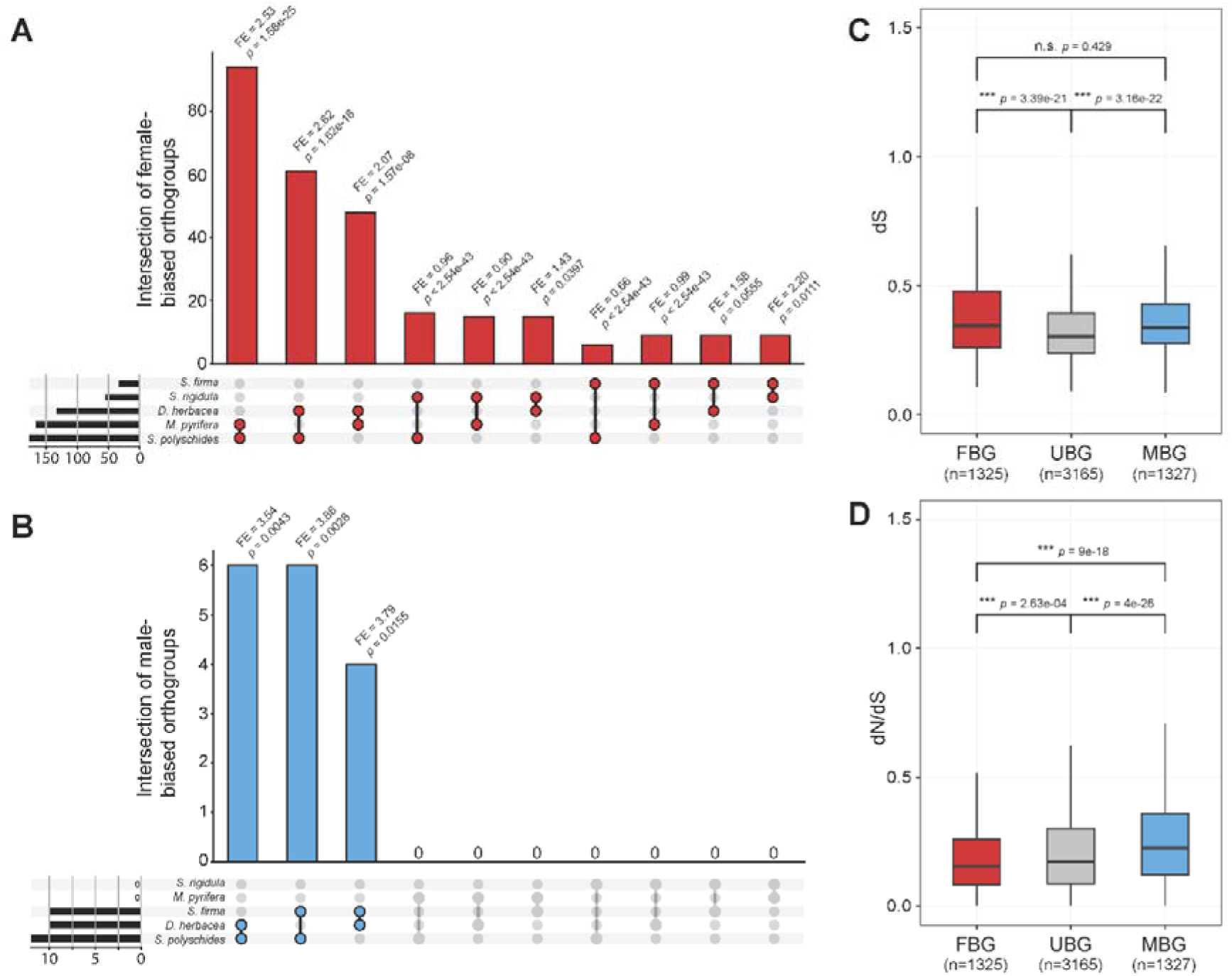
Turn-over of sex-biased gene expression of *Macrocystis pyrifera* gametophytes. (A,B) Intersection plots showing the occurrence of sex-biased orthologs among the five brown algae species *Desmarestia herbacea*, *M. pyrifera*, *Saccorhiza polyschides*, *Sphacelaria rigidula*, and *Sphaerotrichia firma*, based on (A) female-biased genes and (B) male-biased genes in *M. pyrifera* (considering sex bias in at least one population); fold enrichment (FE) and *p*-values are obtained by multi-set intersection analysis. (C,D) Sequence divergence measured as (C) synonymous substitution rate (dS) and (D) ratio of non-synonymous over synonymous substitutions (dN/dS) of sex-biased (FBG, female-biased genes; MBG, male-biased genes) and unbiased genes (UBG) that are orthologous between *M. pyrifera* and *Saccharina japonica* (Wilcoxon test; note that outliers are omitted for clarity). Boxplots depict the median as a horizontal line, interquartile range (IQR) as a box, and a range extended by 1.5*IQR as whiskers.

In addition, while both FBG and MBG have accumulated significantly more synonymous substitutions (dS) than unbiased genes (Figure 2C; Wilcoxon test on orthologous genes between *M. pyrifera* and *Saccharina japonica* ^36^), the ratio of nonsynonymous over synonymous substitutions (dN/dS) was significantly lower in FBG compared to unbiased genes, and significantly higher in MBG (Figure 2D). Therefore, it appears that MBG are more species-specific and present faster evolutionary rates than FBG. It is possible that sexual selection, acting mainly on males, is driving the evolution of male-biased gene expression patterns. A similar pattern was observed in the brown algae *S. polyschides* and *Sphacelaria rigidula*, which display oogamy and anisogamy, respectively, but not in the oogamous *D. herbacea*^34^. Note that increased evolutionary rates of MBG between species have been largely documented in animals (reviewed by Grath & Parsch^37^) while similar patterns have not been found in plants^38–40^. Remarkably, MBG of brown algae with XX/XY systems in the order Fucales do not experience high turnover rates^41^, which is potentially related to the young age of their XX/XY system (65 Mya)^23,42^, compared to the very old U/V system (> 450 Mya)^23,42^. Thus, it appears that patterns of sequence evolution in sex-biased genes vary significantly across species of brown algae. This species-specific pattern of gene evolution may reflect positive selection pressures driving the high turnover of genes involved in sex bias across brown algae, the diversity of gamete dimorphism and the types of life cycles^43^.

### Feminisation and de-masculinisation of gene expression underlie the variant phenotype

We next examined how sex-biased gene expression patterns were modified in the variant lines. Consistent with a phenotypic feminisation, the mean expression levels of FBG were significantly higher in the variants than in the WT males (Wilcoxon test; *p*<0.001), but significantly lower than those of WT females (*p*<0.001; Figure 3)**A**. Equivalently, the mean expression level of MBG in the variants was significantly lower in the variants than in the males (*p*<0.0001), but still significantly higher than in females (*p*<0.0001; Figure 3B). We also assessed the expression of sex-biased genes in the variants within each population. A clear trend of transcriptome feminisation (i.e., up-regulation of FBG) and de-masculinisation (i.e. down-regulation of MBG) emerged in the variants when comparing the total number of differentially expressed genes in the variants to the sex-biased genes within their respective populations (Figure 3C-D). The majority of significantly up-regulated genes in the variants was also female-biased in the same population (multi-set intersection analysis, *p*<0.0001; Figure 3C), and the majority of significantly down-regulated genes in the variants was male-biased (*p*<0.0001; Figure 3D). Additionally, the magnitude of differential expression (log_2_FC) for DEG in several comparisons of variant vs. WT males was highly significantly correlated (**Figure S2F**). Together, these observations indicate a convergence to similar core patterns of transcriptome feminisation and de-masculinisation. We cannot, however, fully exclude additional pleiotropic non-sex-related effects of the underlying changes that caused the partial sex reversals.

**Figure 3.**
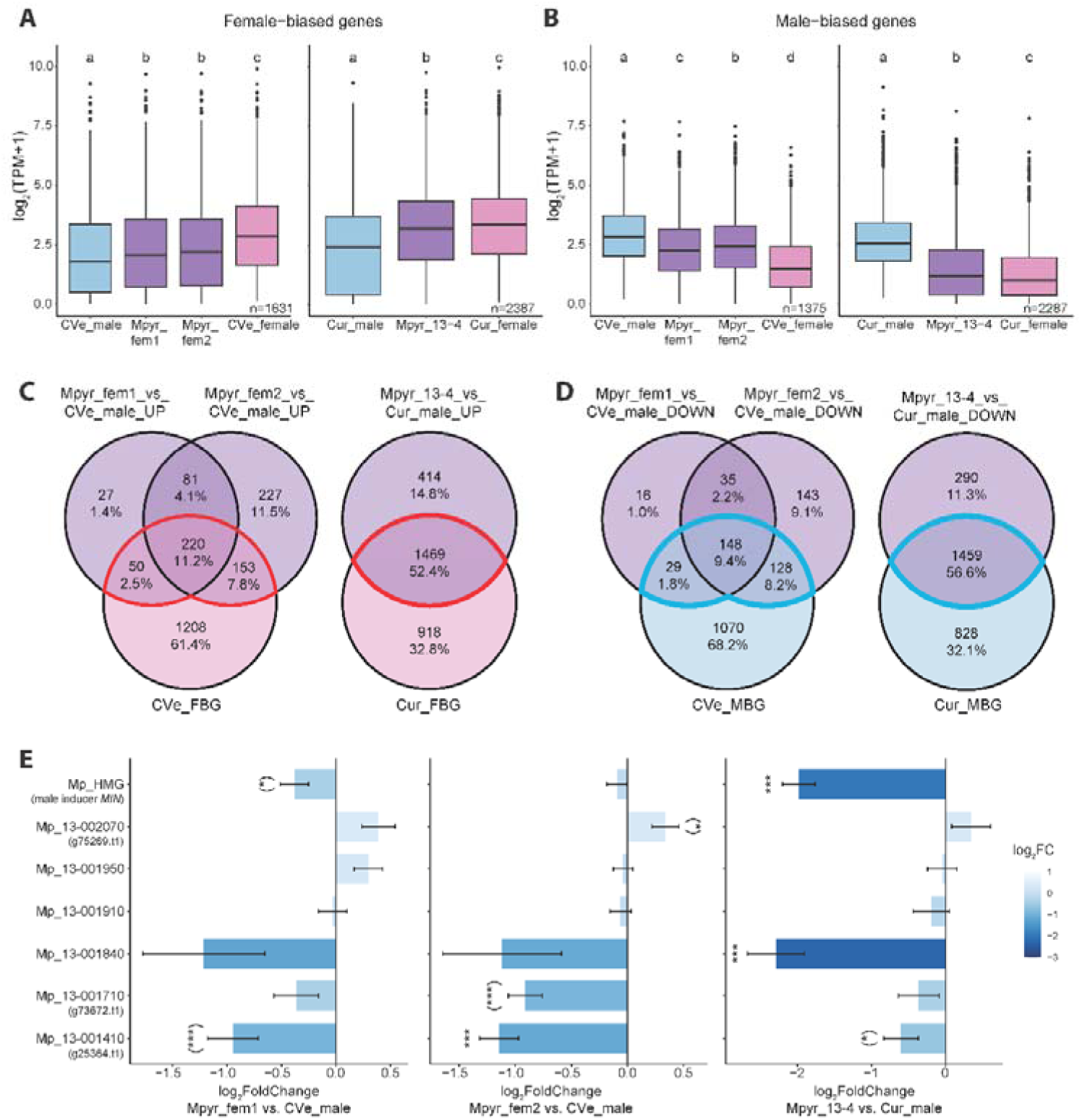
Gene expression patterns of feminisation and de-masculinisation in *Macrocystis pyrifera* variant gametophytes. (A,B). Transcript abundance of (A) female-biased and (B) male-biased genes in *Macrocystis pyrifera* male WTs (CVe_male, Cur_male), feminised male variants (Mpyr_fem1, Mpyr_fem2, Mpyr_13-4) and female WTs (CVe_female, Cur_female). Letters denote statistical differences between strains (Wilcoxon test, *p*<0.01). (C,D) Venn diagrams depicting the absolute number and fraction of significantly (C) up-regulated and (D) down-regulated genes within the *Macrocystis pyrifera* populations CVe (Mpyr_fem) and Cur (Mpyr_13-4) in comparison with the respective (C) female-biased and (D) male-biased genes. Subsets associated with feminisation (up-regulated female-biased genes) and de-masculinisation (down-regulated male-biased genes) are highlighted by red and blue outlines, respectively (DESeq2, *n*=3 for CVe; *n=*2 for Cur). (E) Differential expression of *Macrocystis pyrifera* sex-linked genes for each feminised variant vs. WT male (mean values ± SE, DESeq2, *n*=3 for Mpyr_fem1, Mpyr_fem2; *n*=2 for Mpyr_13-4). Genes are named according to their *Ectocarpus* orthologues (as in Müller et al.^19^); note that gene names for Mp_13-001410, Mp_13-001710 and Mp_13-002070 are g25364, g73672 and g75269, respectively, in the genome annotation^53^; and Mp_HMG is male inducer *MIN*^20^. Asterisks indicate significance of differential expression (*, *p*<0.05; ***, *p* <0.001), and are reported in parentheses if the expression magnitude threshold of |log_2_FoldChange|>1 is not met.

To identify the potential functional gene categories involved in the feminised phenotype of the variants, we then explored enriched gene ontology (GO) terms among gene sets related to feminisation and de-masculinisation. We extracted the FBG that were significantly up-regulated in the variants (Figure 3C) and the MBG that were down-regulated in the variants (Figure 3D). We then identified GO terms which were significantly enriched (Fisher’s exact test, p<0.05) in all three sex-biased subsets of variant DEG, reasoning that these consistently correlate with sexual differentiation. Among these, we explored individual functional gene annotations to infer the affected pathways more specifically. This approach revealed several shared enriched GO terms associated with feminisation that had predicted functions related to polysaccharide metabolism (**Figure S3, Table**);**S**m**4** ainly affecting mannuronan C-5-epimerases, which indicates a distinct cell wall composition in females and feminised males compared to WT males. On the other hand, transmembrane receptor kinases, leucine rich repeat proteins and serine/threonine kinases were consistently down-regulated in the feminised variants, suggesting a role in de-masculinisation (**Figure S3, Table S4**).

In summary, the partially sex-reversed phenotype of the variants co-occurs with significant feminisation and de-masculinisation of global gene expression patterns. Core shared effector gene functions associated with feminisation and de-masculinisation suggest that protein signalling pathways are involved in sexual differentiation in brown algae, and that polysaccharide and cell wall metabolism are strongly affected by the distinct sexual developmental program of these organisms.

### Genes located on the sex chromosome may play a key role in feminisation

We examined the changes in expression of genes located on the V chromosome of *M. pyrifera* during feminisation. Among seven previously identified ancestral male-linked genes^19,21^ (Figure 3E), three showed a consistently reduced transcript abundance across the variants. Two genes had been identified previously in Mpyr_13-4^19^, and belong to an ancestral set of genes that have been consistently male-linked during brown algal evolution^21,23^: the male-master determining gene *Mp_HMG* (male inducer *MIN*)^17,20^, and *Mp_13-001840*, a transmembrane protein (Figure 3E). In addition, both variants Mpyr_fem1 and Mpyr_fem2 show significant down-regulation of *Mp_13-001410*, which encodes a protein with a WD40/YVTN repeat-like-containing domain^44^. This gene was recently integrated into the SDR of kelps^21^ and may therefore have taken on a role in male sexual differentiation.

We used RNA-seq read mapping data for sequence variant calling, followed by manual curation of candidate loci using JBrowse^45^, and did not identify any sequence variation on coding sequences of SDR genes. This suggests that the repression of male SDR genes described here may be controlled by autosomal factors or epigenetic regulators, acting upstream of the SDR and controlling both the activation of male-linked (V-SDR) genes and the repression of the feminised background.

Our dataset comparison further allowed us to approximate a “core” female-restricted gene set that is presumably crucial for the establishment of a functional female program. We identified 24 genes which were exclusively expressed by the WT females within each population (TPM>5^th^ percentile) and additionally differentially higher expressed in WT females than in all genetic males including the variants (**Table S2**). This gene set is enriched in putative functions related to the GO term “nucleotide binding” (Fisher’s exact test, *p*=8.2*10^−12^; **Table S5**). Remarkably, protein sequence alignments using blastp revealed that more than half (14 out of 24) of these are genes located on the sex chromosome of *M. pyrifera* (scaffold_2 in female reference genome by Diesel et al.^31^; **Table S5**). Additionally, seven of these have orthologs on the sex chromosome of *Ectocarpus* (chromosome 13; aligned using the ORCAE database^44^), including two genes which have been shown to be sex-linked in *M. pyrifera* ^21^. Gene *g33225* has strong sequence similarity to *Ectocarpus* gene *Ec-sdr_f_000170* which is a conserved female sex-linked gene across brown algae taxa. In this case, it is likely a gametolog to *Mp_13-001840* described above^21,23^, making both genes of this gametolog pair candidates for genes involved in important steps of sex-determination. Similarly, the ortholog of gene *g35582* is the conserved female-linked *Ectocarpus* gene *Ec-sdr_f_000010,* which is likely a gametolog to the male-linked *Mp_13-001910.* The female gametolog codes for a STE20 protein kinase and has been suggested as central to female development in *M. pyrifera* ^19^ and across brown algae^23^. Note that we cannot exclude that some or all of these genes are located in the sex determining region of *M. pyrifera*, as the SDR boundaries of this species have not been determined yet.

Together, our results suggest that the repression of a combination of male SDR genes is associated with the loss of the male developmental phenotype. Therefore, the role of *MIN* as master sex determinant may be complemented by other auxiliary sex-differentiation factors encoded by the male SDR. At the same time, we identified a core set of female-restricted genes, including genes on the female SDR, which are presumably crucial for the expression of functional female phenotypes.

### Enrichment of regulatory genes underlies male-specific development

In addition to the differential gene expression analysis, we also performed a weighted gene co-expression network analysis (WGCNA)^46^ to identify modules of co-regulated genes and to relate these to the male and female developmental patterns across all samples simultaneously. We assigned four traits to each replicate sample: mean gametophyte cell size (Figure 1B), mean gamete cell size (Figure 1C), genetic sex (1=U-chromosome, 0=V-chromosome), and parthenogenesis (1=present, 0=absent). After removing genes with null expression, we clustered 20,054 genes into 39 modules with a minimum size of 30 genes per module, which are named after colours by default (Figure 4). Size of the modules ranged from 33 genes in module magenta4 to 5075 genes in module blue.

**Figure 4.**
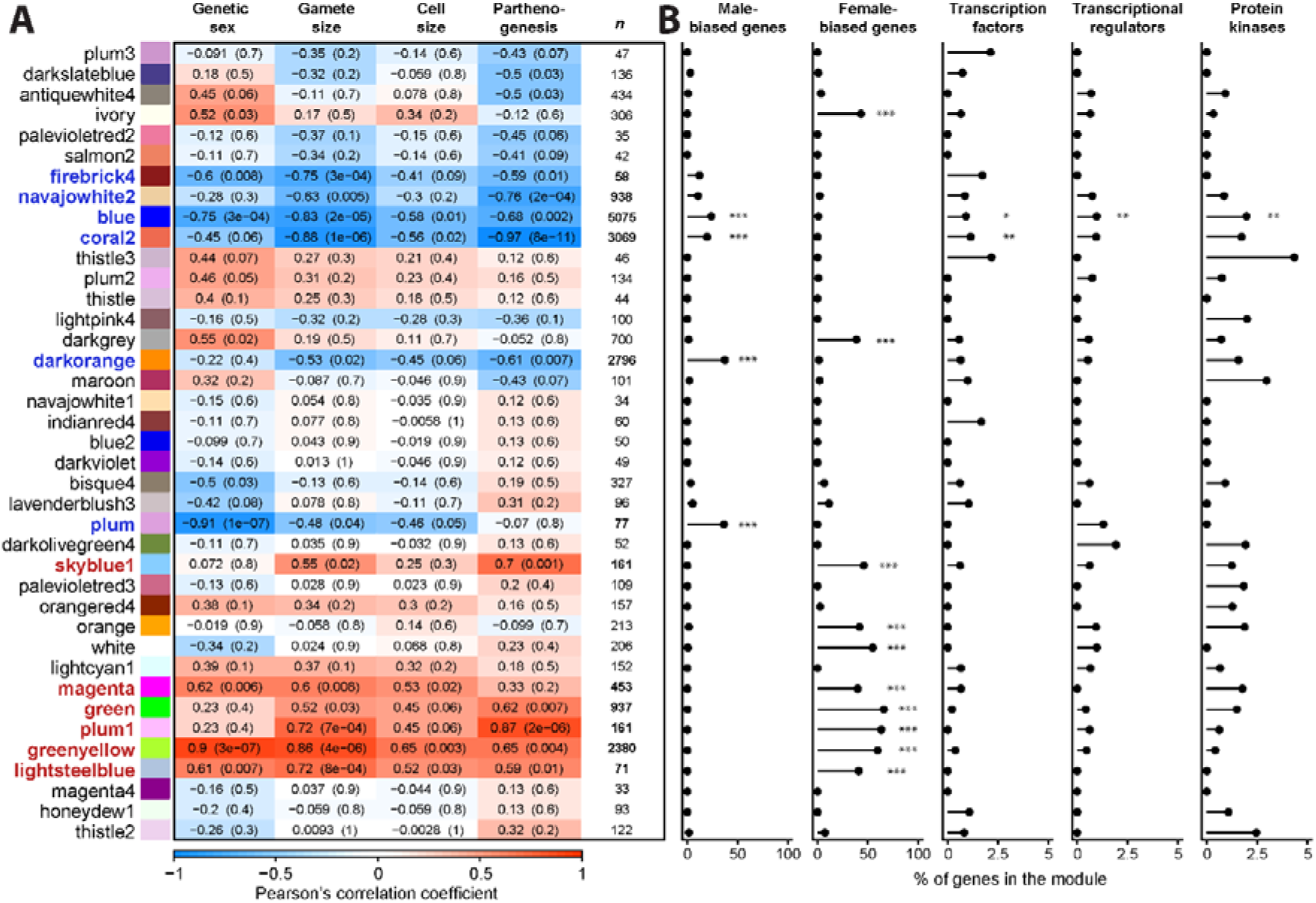
Properties of the 39 modules identified in the weighted gene co-expression network analysis of *Macrocystis pyrifera* WT males, feminised males and WT females. (A) Relation of module eigengenes to the assigned traits genetic sex, gamete size, gametophyte cell size, and parthenogenetic capacity. For each module and trait, Pearson’s correlation coefficient (R) and *p*-value in parentheses are given. Twelve modules with |R|≥0.6 and *p*-value≤0.01 for at least one trait are highlighted in bold and blue (male trait-related) or red (female trait-related) font. Number of genes per module (*n*) is given on the right. (B) Prevalence of sex-biased genes and genes with regulatory function per module, with significant enrichment indicated by asterisks (fold-enrichment>1 for all; ***, *p*<0.0001; **, *p*<0.001; *, *p*<0.05; multi-set intersection analyses).

We identified twelve modules related to sex phenotype differentiation, which showed a significant Pearson’s correlation (|R|≥0.6; *p*≤0.01) to at least one trait (highlighted in bold font in Figure 4A). Of these, six modules were negatively correlated with female traits (blue, coral2, navajowhite2, plum, firebrick4, darkorange) and six modules were positively correlated with female traits (greenyellow, plum1, skyblue1, lightsteelblue, magenta, green). Accordingly, most of these modules were significantly enriched (multi-set intersection analysis) with genes that we had previously identified as sex-biased (Figure 4B). Notably, the three largest modules, blue, coral2 and darkorange, were strongly negatively correlated to traits associated with feminisation, and, accordingly, were significantly enriched in male-biased genes (multi-set intersection analysis, *p*<0.0001; Figure 4A**,B**). The six modules related to male traits consisted of in total 12013 genes (54.15% of the whole transcriptome), whereas the six female trait-related modules consisted of 4163 genes (18.76%).

GO term enrichment analysis of these twelve modules highlighted the widespread involvement of cellular components related to chromatin structure and epigenetic histone marks in the male-related modules, e.g. blue, coral2 and darkorange (e.g., “chromatin”, “histone acetyltransferase complex”, “nucleosome”; Fisher’s exact test, *p*<0.0001 for the above modules; **Table S4**). Histone modifications play a major role in the life cycle control of brown algae^47–49^. In the model *Ectocarpus*, genes involved in male-specific development were mostly enriched in activation-associated marks in males compared to females^48^, suggesting a link between chromatin and activation of the male developmental program. Note that the significant enrichment of “cilium movement” in module darkorange (*p*=5.7*10^−21^) is also consistent with male sperm production. In addition, the enrichment of regulatory molecular functions in, e.g., modules blue, coral2 and green (e.g., “protein binding”, “DNA binding”, “RNA binding”; *p*<0.001 for the above modules; **Table S4**) suggests that sexual differentiation relies on mechanisms spanning from regulation of gene expression to posttranslational protein modifications.

Based on the above indication, we annotated motifs related to regulatory protein activity in the genome using the Plant Transcription factor & Protein Kinase Identifier and Classifier (iTAK^50^; Figure 4B; **Table S2**). Remarkably, the two largest male-trait related modules were significantly enriched in regulatory gene functions. Module blue was significantly enriched in transcription factors (0.91%; multi-set interaction analysis, *p*=0.049), transcriptional regulators (0.97%; *p*=0.010) and protein kinases (1.97%; *p*=0.004), while module coral2 was significantly enriched in transcription factors (1.14%; *p*=0.004). In comparison, the largest female trait-related modules green and greenyellow were highly significantly enriched in GO terms of metabolic functions related to, e.g., pigment synthesis (Fisher’s exact test, *p*=6.8*10^−20^) and cell wall composition (Fisher’s exact test, *p*=8.5*10^−11^; **Table S4**).

Together, and in combination with the trend of larger male trait-related gene co-expression modules, this analysis suggests that male development requires substantial restructuring of the cellular machinery involving mechanisms from chromatin and epigenetic remodelling to posttranslational modifications, in comparison to the female developmental program, which targets metabolic processes more directly.

### A female-like phenotype is the “ground state” of kelp gametophyte morphology

We show that the three male *M. pyrifera* variants Mpyr_13-4^19^, Mpyr_fem1 and Mpyr_fem2 (this study), which were obtained in independent experiments from two distinct WT male gametophytes, present a similarly feminised phenotype. Despite being genetically male, they all share typical female features such as increased gametophyte cell and gamete dimensions, and parthenogenetic capacity. However, the size of the gametophyte and gamete cells does not fully reproduce a female phenotype, and the gametes are sterile. Therefore, they are partially sex-reversed, displaying an intermediate phenotype to WT males and females.

These phenotypes occurred despite the presence of the male V-SDR and the absence of the female U-SDR. The fact that the transcriptome of the variants was significantly enriched in FBG demonstrates that these are not under direct control of the female sex locus, but rather that their expression may be repressed in WT male gametophytes, suggesting dominance of the male (V) over the female (U) sex locus. In the model brown alga *Ectocarpus*, the V haplotype is dominant over the U haplotype^49,51,52^, implying that the male developmental program triggered in the presence of a V-SDR would be superimposed on the background of a “default” female development^17,49^. Additionally, transitions to hermaphroditism across brown algae orders have consistently arisen from a male background via introgression of female SDR genes^23^. This suggests that, at least in this lineage, haploid males may more easily gain female traits than vice versa by acquiring few genes crucial for female function^23^.

We therefore propose that a female-like phenotype is the “ground state” of giant kelp gametophyte morphology. The initiation of the male developmental program requires the presence of a V-SDR that includes the master male sex determining gene Mp_HMG (male inducer *MIN*)^20,49^ together with accessory V-SDR linked genes, which coordinate male development and the associated transcriptomic changes^20,49^. The male V-SDR therefore operates by repressing the female-like ground state while activating the male developmental program. The female U-SDR appears to be crucial to the production of functional eggs, with the expression of a core set of U-linked genes providing key functions to complement the ground state developmental program and producing a functional female.

## Acknowledgments

We thank Dieter Müller for providing the strains used in this study, and Martin Gachenot for support in the ploidy analysis. We thank the BMBF-funded de.NBI Cloud within the German Network for Bioinformatics Infrastructure (de.NBI) (031A532B, 031A533A, 031A533B, 031A534A, 031A535A, 031A537A, 031A537B, 031A537C, 031A537D, 031A538A). Computations were also performed in the MPCDF Cobra supercomputer and the cluster of the Max Planck Campus in Tübingen, Germany. This work was supported by the MPG, the ERC (grant n. 864038 to SMC) the Moore Foundation (GBMF11489, SMC) and the Bettencourt-Schuller Foundation (SMC). JBR was supported by a Humboldt Research Fellowship for postdoctoral researchers from the Alexander von Humboldt Foundation.

## Author contributions

DL: Investigation (lead); Formal analysis (lead); Visualization (lead), Writing – original draft (lead), Writing – review and editing (supporting).

MZ, OG, JBR: Investigation (equal); Methodology (supporting)

GC: Investigation (equal), Methodology (equal); Visualization (supporting)

FBH: Data curation (lead); Visualization (supporting)

SMC: Conceptualization (lead); Funding acquisition (lead); Methodology (equal); Project administration (lead); Visualization (supporting); Writing – original draft (supporting); Writing – review and editing (lead).

## Supplemental information

**Table S1** List of *Macrocystis pyrifera* strains used and genome and transcriptome assembly statistics, related to STAR Methods.

**Table S2** Expression data and functional annotation of all genes of *Macrocystis pyrifera*, related to STAR Methods, Figures 3-4.

**Table S3** List of single-copy orthologs across the five brown algae species *Macrocystis pyrifera*, *Sphacelaria rigidula*, *Sphaerotrichia firma*, *Desmarestia herbacea*, *Saccorhiza polyschides* (Cossard et al.^34^), related to Figure 2.

**Table S4** Significantly enriched gene ontology (GO) terms in *Macrocystis pyrifera* gametophytes, related to Figures 3-4.

**Table S5** Alignment results and gene ontology (GO) enrichment results of female-restricted genes of *Macrocystis pyrifera* gametophytes to reference genomes of *Macrocystis pyrifera* and *Ectocarpus* species 7, related to Figure 3.

**Figure S1** Information on the *Macrocystis pyrifera* strains used in this study, related to STAR Methods, Figure 1.

**Figure S2** Dynamics of sex-biased gene expression of *Macrocystis pyrifera* WT and variant gametophytes, related to Figures 2-3.

**Figure S3** Number of genes within each significantly enriched gene ontology (GO) term shared by all within-population comparisons of feminised males and females vs. WT males of *Macrocystis pyrifera*, related to Figure 3.

## STAR Methods

### Resource availability

#### Lead contact

Further information and requests for resources and reagents should be directed to and will be fulfilled by the lead contact, Susana M. Coelho.

#### Materials availability

This study did not generate new unique reagents.

#### Data and code availability

- The RNAseq and DNAseq data generated for this study have been deposited at NCBI (BioProject PRJNA1072278). The modified *M. pyrifera* reference genome and annotation files have been deposited at EDMOND (https://doi.org/10.17617/3.G6SL3X). The additional RNAseq dataset by Müller et al.^19^ and by Cossard et al.^34^ are available at NCBI. Accession numbers and DOIs are listed in the key resources table.
- This paper does not report original code.
- Any additional information required to reanalyze the data reported in this paper is available from the lead contact upon request.

### Experimental model and study participant details

Fertile sporophytes of the giant kelp *Macrocystis pyrifera* (RRID:NCBITaxon_35122) were collected at Curaco de Vélez, Chiloé, Chile, in 2006 (**Figure S1**). Meiospores were released and clonal cultures were established of two unialgal gametophyte isolates CVe13f female (CVe_female) and CVe30m male (CVe_male). The male gametophyte strain CVe30m was treated with colchicine as described by Müller et al.^19^, to obtain individuals with feminised phenotypes. In short, gametophytes were grown on agar plates (1% agar in seawater) in contact with a 6 mm filter paper loaded with 1 mg of Colchicine (Fluka, Honeywell Research Chemicals, Illkirch, France). After 12-16 weeks of culture (12 ± 2°C, 2-3 µmol photons m-2 s-1 from daylight type fluorescent lamps under a 14:10 h light:dark cycle), the feminised variants Mpyr_fem1 and Mpyr_fem2 were isolated from a phenotypically heterogeneous gametophyte regenerate. The mechanism by which colchicine induces the feminised phenotype is as yet unknown. As colchicine acts by inhibiting tubulin assembly and suppressing microtubule formation, the treatment may have induced chromosome segregation problems during mitosis and potentially sequence duplications or deletions.

All cultures were then maintained vegetatively under 25-30 µmol phot. m^-2^ s^-1^ red LED light^72^ (MaxLED 500 RGBW, Paulmann, Springe, Germany) in a thermostatic cabinet (TC 445 L, Lovibond, Dortmund, Germany) in a 14:10 h light:dark cycle at 14°C in 50% Provasoli-enriched natural seawater (PES^73^ with iodine enrichment^74^).

### Method details

#### Phenotypic characterization

To conduct measurements of cell and gamete size, gametophyte tufts were ground carefully in an Eppendorf tube using a micropestle. Gametophyte fragments were sieved to obtain a fraction smaller than 100 µm and settled in 12 mL 100% PES in petri dishes. For measurements of vegetative cell size, gametophytes were grown on coverslips within petri dishes for two weeks under red light. To increase contrast, cells were stained with a droplet of lactophenol blue on the coverslips, which were then inverted onto a glass slide and images were taken at 400x magnification with a microscope camera (acA2440-75um, Basler, Ahrensburg, Germany) fitted to an inverted microscope (Axio Vert.A1, Zeiss, Jena, Germany). All measurements were conducted with ImageJ Fiji^56^.

Fertilization of ground gametophyte fragments was induced after 2 days at reduced irradiance of 5-10 µmol photons m^-2^ s^-1^ white LED light in a 14:10 h light:dark cycle (aquaLUMix AMAZON-GROW, LEDaquaristik GmbH, Hövelhof, Germany) at 12°C (TC 135 S, Lovibond, Dortmund, Germany) to facilitate gametophyte recovery from grinding. To induce fertility, irradiance was raised to 20-30 µmol photons m^−2^ s^−1^. After 7 days, dishes with Mpyr_fem1 were moved to 50-70 µmol m^−2^ s^−1^ as no fertility had been observed until then. Medium was changed and cultures were observed every 3-4 days. Once the cultures were undergoing gametogenesis, photos were taken to measure gamete size and experimental crosses were produced by addition of fertile male gametophytes to determine whether gametes of the feminised strains were capable of syngamy.

Genetic sex of all gametophytes was confirmed using PCR-based markers (see key resources table; Lipinska et al.^54^) with the following conditions: initial denaturation for 3min at 95°C; followed by 35 cycles of denaturation for 30s at 95°C, annealing for 30s at 65°C decreasing to 55°C by 1°C/cycle for the first ten cycles, elongation for 30s at 72°C; and final elongation for 10min at 72°C. For the determination of ploidy, small tufts of gametophytes (20-50 mg fresh weight) were patted dry and cut to fine fragments in 500 µL of nuclei isolation buffer^75^ (30 mM MgCl2, 20 mM sodium citrate, 120 mM sorbitol, 55 mM HEPES, 5 mM EDTA, 0.1% (v/v) Triton X-100, pH 7.5), with addition of 0.5 µg/L Proteinase K, 5‰ SDS, 0.1 mM Tris HCl pH 8.0. Nuclei were strained through a 10 µm nylon mesh and stained using 1X SYBR Green I (Invitrogen, Waltham, USA). After 10 min incubation, DNA content was measured in a flow cytometer (NovoCyte Advanteon, Agilent, Santa Clara, USA).

#### DNA extraction and sequencing

Genomic DNA was isolated from 5-15 mg blotted dry gametophytes using the OmniPrep Genomic DNA Isolation kit (G-Biosciences, St. Louis, USA) following the manufacturer’s instructions. Libraries were prepared using a custom protocol for Tn5 tagmentation (Tn5 expressed and purified according to Picelli et al.^76^). 10 ng of DNA were tagmented in 20 µL TAPS-DMF buffer for 7 min at 55°C. Tn5 was stripped from DNA by adding 2 µL 1% SDS and inactivated by incubating at 70°C for 10 min. Tagmented DNA was purified using a DNA Clean and Concentrator 5 Kit (Zymo Research, Irvine, USA) before PCR amplification using Q5 High-Fidelity DNA Polymerase (New England Biolabs, Ipswich, USA) with the following conditions: 72°C for 5min, 98°C for 30s, and 8 cycles of 98°C for 15s, 67°C for 30s, 72°C for 60s. Size selection of PCR-enriched samples was performed using AMPure XP beads (Beckman Coulter, Pasadena, USA) to retain fragments between 300-550 bp. DNA purity was confirmed by spectrophotometry (BioSpectrometer, Eppendorf, Hamburg, Germany), concentrations were measured by fluorometry (dsDNA High Sensitivity Assay, Qubit 4, Invitrogen, Waltham, USA) and library size distribution was confirmed using capillary electrophoresis (High Sensitivity DNA Assay, Agilent 2100 Bioanalyzer, Agilent Technologies, Santa Clara, USA). DNA was sequenced as 150 bp paired end libraries to an average depth of 39–59x on an Illumina HiSeq3000 (CVe_male, Mpyr_fem1) or NextSeq2000 (Mpyr_fem2) at the Genome Core Facility at the MPI Tübingen Campus.

#### Coverage analysis to identify genomic duplications

Following quality trimming of raw reads with Trimmomatic v.0.39^77^ and PCR duplicate removal using Picard MarkDuplicates v.2.27.4 (http://broadinstitute.github.io/picard/), reads were mapped against the *Macrocystis pyrifera* genome published by Diesel et al.^31^ using BWA-MEM v.0.7.17^58^. To identify chromosome duplications or deletions in the variant strains, we compared normalised sequence coverage to the chromosome-level scaffolds between variant strains and the wild type strain. Per-base coverage was calculated using genomecov within BEDTools v.2.27.1^78^, from which average coverage per scaffold and per genome were calculated. Average scaffold coverage was then normalised by whole-genome coverage, yielding values fluctuating around 1. Genomic regions with normalised coverage <0.5 were considered weakly covered, whereas regions with normalised coverage >2 were considered overrepresented. Applying a filter extracting scaffolds that were averagely covered in the wild type male (0.5 ≤ normalised coverage ≤ 2) and had a strong coverage bias between wild type and feminised variant (difference ≥ 0.5) yielded no evidence for duplications or deletions in the variants.

#### RNA extraction and sequencing

Colonies of WT CVe_male and CVe_female cultures, and the feminised male variants Mpyr_fem1 and Mpyr_fem2 were fragmented using tweezers and cultivated each in four replicate petri dishes (IZJ 55 mm) filled with 12 mL 100% PES at fertility-inducing conditions (see above) for 7 days. At the time of collection, reproductive features were not yet visible (to be comparable with Müller et al.^19^). Four lentil-sized replicates (approx. 20 mg fresh weight) per strain were patted dry in tissue paper, snap-frozen in liquid nitrogen and stored at −80°C.

Frozen biomass was homogenised in 1.5 mL centrifuge tubes using a micropestle while preventing thawing of the sample by submerging the tube frequently in liquid nitrogen. Lysis was performed in 1 mL CTAB3 buffer (100 mM Tris-HCl pH 8.0; 1.4 M NaCl; 20 mM EDTA pH 8.0; 2% Plant RNA Isolation Aid [Invitrogen, Waltham, USA]; 2% cetyltrimethyl ammonium bromide; 1% β-mercaptoethanol) for 15 min at 45°C. The organic phase was separated twice using the same volume of 24:1 chloroform:isoamylalcohol and centrifuging at 10,000 g for 15 min at 4°C. The upper aqueous phase was collected and RNA was precipitated overnight at −20°C in a solution of 3 M LiCl and 1% β-mercaptoethanol. RNA was collected by centrifuging at 18,200 g for 60 min at 4°C and the supernatant was discarded. The RNA pellet was washed twice in 1 mL 80% ice-cold Ethanol, air-dried and eluted in 30 µL nuclease-free water. DNA was removed using TURBO DNase (Invitrogen, Waltham, USA) according to the manufacturer’s instructions and 35 µL of the supernatant were treated with the RNA Clean & Concentrator Kit 5 (Zymo Research, Irvine, USA) using an adapted protocol (the RNA prep buffer step was performed twice; RNA was washed in 700 µL wash buffer four times) and RNA was eluted in 15 µL nuclease-free water. RNA purity was confirmed by spectrophotometry (BioSpectrometer, Eppendorf, Hamburg, Germany), concentrations were measured by fluorometry (RNA Broad Range Assay, Qubit 4, Invitrogen, Waltham, USA) and RNA size distribution was confirmed using capillary electrophoresis (Plant RNA Nano Assay, Agilent 2100 Bioanalyzer, Agilent Technologies, Santa Clara, USA). Sequencing libraries were prepared using a commercial kit (NEBNext Ultra II Directional RNA Library Prep with Sample Purification Beads, New England Biolabs, Ipswich, USA), and cDNA of three replicates per strain was sequenced as 150 bp (100 bp for Mpyr_CVe13f) paired end libraries on an Illumina NextSeq2000 at the Genome Core Facility at the MPI Tübingen Campus.

#### Transcriptome analysis

Additionally to the data produced in this experiment, we re-analysed an RNA-seq dataset including the colchicine-treated feminised male *M. pyrifera* gametophyte Mpyr_13-4 and the two WTs Cur6m male (Cur_male) and Cur4f female (Cur_female), which had been produced in an independent experiment using a different male gametophyte^19^. This yielded a dataset with three feminised variants, two WT males and two WT females from two independent experiments.

Read quality and adapter trimming, mapping and gene expression values were performed according to the pipeline by Perroud et al.^79^, using gmap-gsnap version 2021-12-17^60^ and subread version 2.0.3^61^. The *M. pyrifera* genome by Lipinska et al.^53^ was used as reference and modified on the SDR contigs. Differential expression between samples within the two independent experiments was assessed using DESeq2 version 1.38.2^33^ (see below). Transcript abundances were calculated as transcript per million (TPM) and normalised as log(TPM+1). For each sequencing library, all genes with TPM values > 5^th^ percentile, excluding genes with null expression, were considered as expressed.

We inferred gene orthology between the five brown algae species *M. pyrifera*, *Desmarestia herbacea*, *Saccorhiza polyschides*, *Sphacelaria rigidula* and *Sphaerotrichia firma* using Orthofinder v.2.5.2^66^, accessing the genomes published by Cossard et al.^34^. All orthologous genes were ‘expressed’ according to the above definition (TPM > 5^th^ percentile)^34^. Enrichment of sex-biased genes among all orthologs was analysed using multi-set intersection analysis (see below). To test for differences in rates of evolutionary divergence between different categories of sex-biased and unbiased genes, single-copy orthologous genes were identified for *M. pyrifera* and *Saccharina japonica*^36^ using Orthofinder with default parameters. Orthologous proteins between species pairs were aligned with MAFFT v.7.453^67^, and the alignments were curated with Gblocks v.0.91^68^ and back-translated to nucleotides using translatorX^69^. We used these nucleotide alignments as input for phylogenetic analysis by maximum likelihood (CodeML in PAML v.4.9^70^) to infer pairwise dN/dS with the F3×4 model of codon frequencies.

Weighted gene co-expression network analysis (WGCNA) was then conducted on the concatenated dataset using R package WGCNA^46^. To account for baseline differences between the two experiments, batch effects were removed from the log(TPM+1)-normalised data using removeBatchEffect in limma^80^. A signed network was constructed using a biweight midcorrelation (“bicor”), a soft thresholding power of 18, and maximum portion of outliers of 0.05. To obtain a reasonable number of gene co-expression modules with sufficient size for statistical analyses, modules were constructed with a minimum size of 30 genes and were merged to a correlation threshold of 0.75. We correlated gene co-expression to the traits genetic sex (1=female, 0=male), mean gamete size, mean cell size, and parthenogenesis (1=present, 0=absent). Modules with strong Pearson’s correlation (|R|≥0.6) and significance (*p*≤0.01) to one or several traits were identified for further analysis.

Functional annotation of genome protein sequences against the NCBI non-redundant (NR) database was performed using Diamond^62^ and only top hits were retained. Of 22186 annotated gene products, 19910 (89.74%) were functionally annotated. Annotation using the Kyoto Encyclopedia of Genes and Genomes (KEGG)^64^ covered 17.38% of genes. Gene ontology (GO) annotation had been performed previously using Blast2GO^19,63^, and covered 10460 genes (47.15%). Among all protein sequences in the genome, 160 transcription factors, 157 transcriptional regulators and 344 protein kinases were annotated using the online tool Plant Transcription factor & Protein Kinase Identifier and Classifier (iTAK)^50^. All functional annotation results are collated in **Table S2**.

Variant calling and filtering was performed according to Haas et al.^81^ with GATK package version 4.2.6.1^65^. Lists of sequence variants were compared between strains to retain only variants specific to WT males and feminised males. All sequence variations appearing only in one dataset (WT male vs. feminised male) were marked as potential candidates which were manually curated in JBrowse^45^.

### Quantification and statistical analysis

All statistical analyses were produced in R version 4.2.1^55^. Number of statistical replicates (*n*) and the applied statistical tests are reported in the Figures and/or in the Figure legends. Asterisks in plots represent p values as follows: *, p<0.05; **, p<0.01; *** p<0.001.

Significant differences between strains in cell and gamete size, and between expression levels of sex-biased genes were analysed by non-parametric Kruskal-Wallis rank sum tests followed by pairwise Wilcoxon rank sum tests with false discovery rate (FDR)^82^ *p*-value adjustment^82^ (padj<0.05).

The significance threshold for differentially expressed genes in DESeq2 was considered at log_2_ fold-change (log_2_FC)>1 and FDR^82^-adjusted padj<0.001. All genes differentially expressed between the respective female and male WTs per experiment (CVe_female vs. CVe_male; Cur_female vs. Cur_male) were considered sex-biased genes. Genes with statistical support for non-differential expression in these comparisons (log_2_FC<1, padj<0.001) were considered unbiased.

Significant enrichment of sex-biased genes among the up- and down-regulated genes of feminised variants in comparison with the WT males, of sex-biased genes among expressed single-copy orthologous genes between species, and of genes with regulatory function within gene co-expression modules was assessed using multi-set intersection analyses (*p*<0.05; SuperExactTest^35^). Differences in evolutionary rates between sex-biased and unbiased genes were assessed with pairwise Wilcoxon rank sum tests (*p*<0.05) after filtering out sequences saturated in synonymous mutations (dS>3).

Significant enrichment of GO terms within differentially expressed gene sets was analysed using Fisher’s exact test (*p*<0.05) within R package topGO^71^. In the GO term enrichment analysis of differentially expressed gene sets related to feminisation and de-masculinisation, only the genes identified as expressed within each population (TPM>5^th^ percentile) were considered as a background data set. As the whole transcriptome was considered in the WGCNA, the entire dataset was used.

