## Supplemental Figures for "Developmental pathways underlying sexual differentiation in a U/V sex chromosome system"

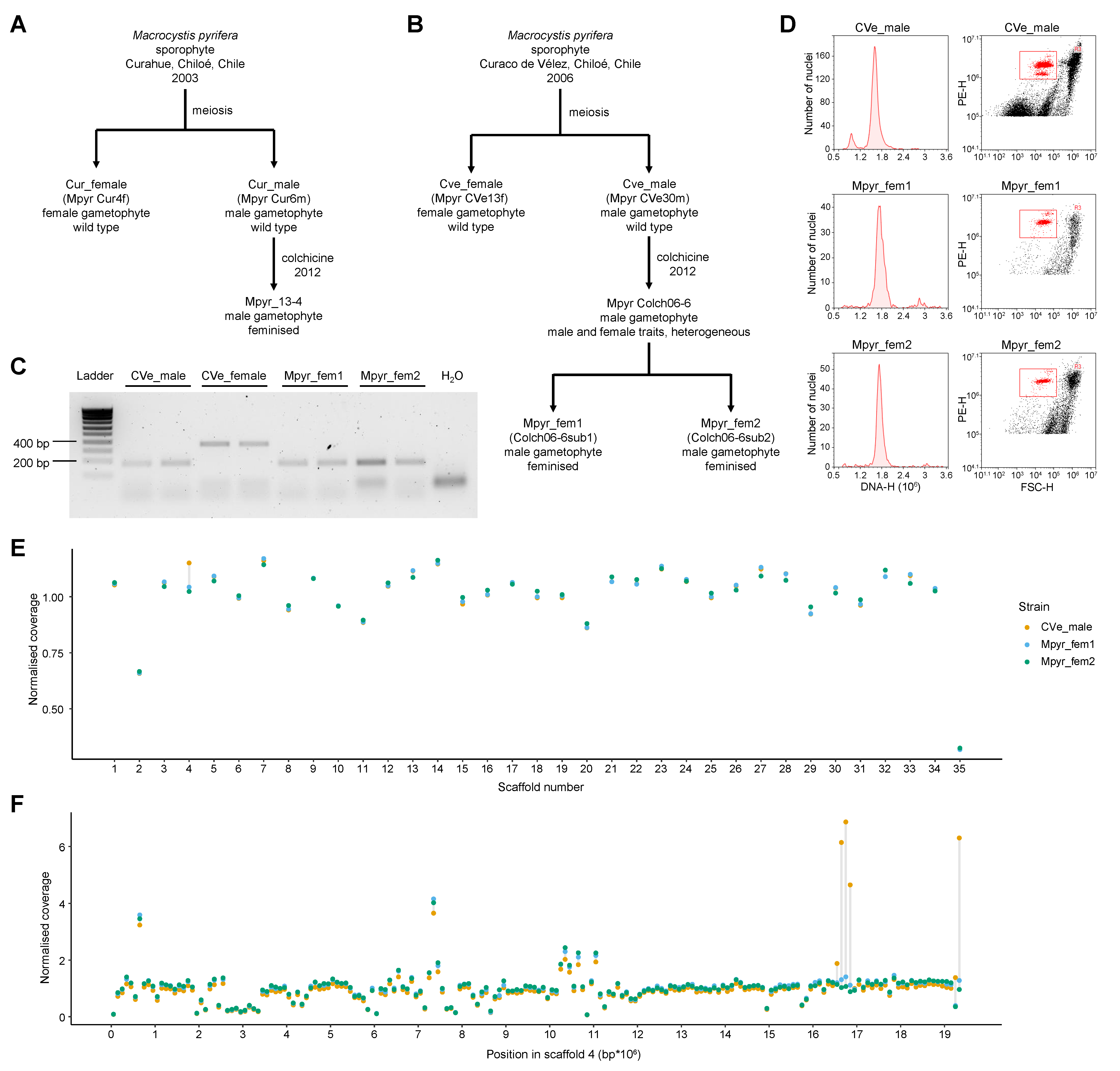

**Figure S1** Information on the *Macrocystis pyrifera* strains used in this study. **(A,B)** Pedigree of strains described **(A)** in Müller et al.^19^ and **(B)** in this study. **(C)** Gel electrophoresis pattern of PCR products of *Macrocysis pyrifera* gametophyte strains CVe_male (wild type male), CVe_female (wild type female), and Mpyr_fem1 and Mpyr_fem2 (feminised males). PCR products were run on a 1.8% Agarose gel for 20 min at 120 V. The target sequence of male marker Mac13.1750.1 (F, TTTCAGGTACGACGTGTTGG; R, CAATTCCGTCCAAGTCGTTT) amplifies at 180 bp, that of female marker M_68_58_2 (F, GTTGGTATAACGGCGTTGGA; R, CACCTCCTTAAAGTTGCGGC; Lipinska et al.^54^) amplifies at 350 bp. PCRs were conducted with both markers present in each reaction. **(D)** Results of flow cytometry analysis to estimate DNA content of *Macrocystis pyrifera* gametophyte strains CVe_male (wild type male), Mpyr_fem1 and Mpyr_fem2 (feminised male variants). Plots in the left column show density plots of fluorescence signals (i.e., number of nuclei) as a function of DNA content (DNA-H); plots in the right column show populations of nuclei grouped by forward scatter (FSC-H) and fluorescence signal (PE-H) in scatterplots. **(E)** Normalised sequencing coverage averaged over each scaffold of the *M. pyrifera* genome^31^. Values represented by coloured dots per strain are connected with a grey line within each scaffold to facilitate comparison. **(F)** Normalised sequence coverage averaged over 100kb windows of scaffold 4 shows that differences in average coverage of scaffold 4 are due to sequence overrepresentation within a ~200kbp region in the wild type male, not due to a deletion in the feminised variants.

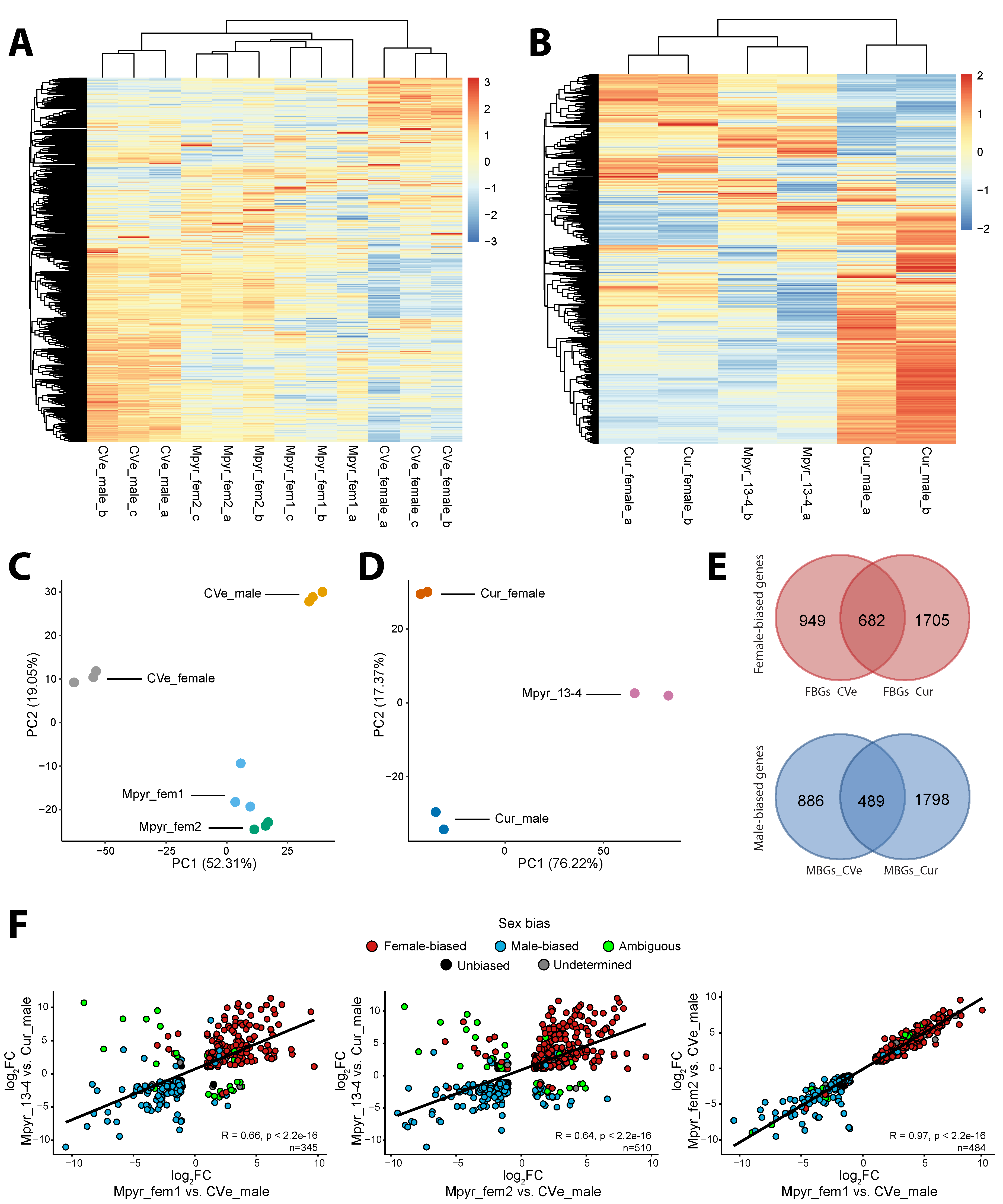

**Figure S2** Dynamics of sex-biased gene expression of *Macrocystis pyrifera* wild type and variant gametophytes. **(A,B)** Global gene expression heatmap and hierarchical clustering based on z-score normalised log(TPM+1) expression values for all expressed genes (TPM > 5^th^ percentile) in male wild types (CVe_male, Cur_male), feminised male variants (Mpyr_fem1, Mpyr_fem2, Mpyr_13-4) and female wild types (CVe_female, Cur_female) for two different experiments (**A**, this study; **B**, Müller et al.^19^). **(C,D)** Principal component analysis (PCA) plots to visualise clustering of sequencing replicates in populations CVe **(C)** and Cur **(D)**, based on normalised expression values for all expressed genes. Percentage of variation explained by each principal component (PC) is indicated. **(E)** Venn diagrams depicting the number of shared sex-biased genes between the populations CVe and Cur based on differentially expressed genes between wild type females and males. **(F)** Correlation of log_2_ fold-change (log_2_FC) between comparisons for differentially expressed genes occurring in two comparisons (co_DEGs) of feminised variants and male wild types. Color indicates sex bias for each gene: red, female-biased in at least one population; blue, male-biased in at least one population; green, male-biased in one population and female-biased in the other (ambiguous); black, unbiased in at least one population and unbiased or undetermined in the other; grey, undetermined in either population.

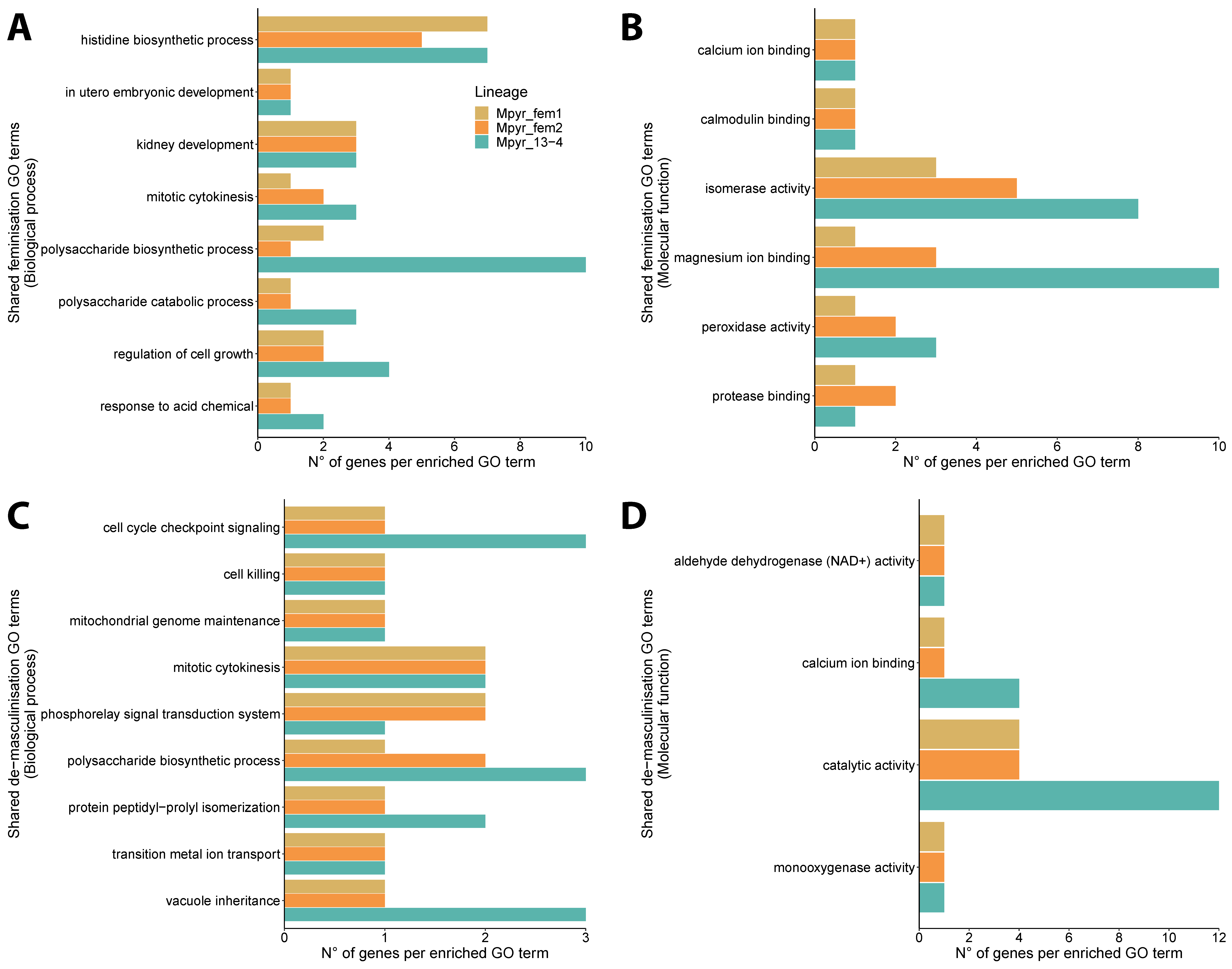

**Figure S3** Number of genes within each significantly enriched gene ontology (GO) term shared by all within-population comparisons of feminised males and females vs. wild type males of *Macrocystis pyrifera*, representing conserved effector gene functions of feminisation **(A,B)** and de-masculinisation **(C,D)** in the GO categories of biological process **(A,C)** and molecular function **(B,D)**.
